# Enterovirus D68 2A protease causes nuclear pore complex dysfunction and independently contributes to motor neuron toxicity

**DOI:** 10.1101/2025.01.23.632178

**Authors:** Katrina M. Zinn, Mathew W. McLaren, Michael T. Imai, Malavika M. Jayaram, Jeffrey D. Rothstein, Matthew J. Elrick

## Abstract

The picornavirus Enterovirus D68 (EV-D68) is an important pathogen associated with acute flaccid myelitis (AFM). The pathogenesis of AFM involves infection of spinal motor neurons and motor neuron death, however the mechanisms linking EV-D68 infection to selective neurotoxicity are not well understood. Dysfunction of the nuclear pore complex (NPC) has been implicated in motor neuron injury in neurodegenerative diseases such as amyotrophic lateral sclerosis, and the NPC is also modified by picornavirus proteases during the course of infection. We therefore sought to determine the impact of EV-D68 proteases on NPC composition and function. We demonstrate widespread disruption of NPC composition by EV-D68 2A and 3C proteases via the direct cleavage of a relatively small number of nucleoporins, notably Nup98 and POM121 by 2A^pro^. Using reporter systems, we demonstrate that 2A^pro^ inhibits nuclear import and export of protein cargoes and also disrupts the permeability barrier of the NPC, while having no apparent effect on RNA export. Independently, we show that 2A^pro^ is toxic to induced pluripotent stem cell derived motor neurons by demonstrating a rescue of toxicity with 2A^pro^ inhibitor telaprevir at concentrations that are insufficient to inhibit viral replication. These findings expand our understanding of EV-D68 neuropathogenesis and provides a rationale for studying the NPC or 2A^pro^ as therapeutic targets in AFM.

## INTRODUCTION

Enterovirus D68 (EV-D68) has caused multiple outbreaks of severe respiratory disease and the polio-like disorder acute flaccid myelitis (AFM) during the past decade. Though much work has been done to demonstrate the association between EV-D68 and AFM and to characterize the molecular evolution of EV-D68 leading to the increased virulence of contemporary strains, the mechanisms underlying its neuropathogenesis remain poorly understood (Elrick et al. 2021). EV-D68 has been shown to infect spinal motor neurons in mouse models (Hixon et al. 2017) and human autopsy tissue (Vogt et al. 2022), and the dysfunction or death of motor neurons is thought to be the proximate cause of paralysis in AFM.

EV-D68 is a member of the family *Picornaviridae*, which have a non-enveloped icosahedral capsid and a single-stranded positive sense RNA genome (Oberste et al. 2004). Picornaviruses alter the composition and function of the nuclear pore complex (NPC) during the course of their life cycle (reviewed in Lizcano-Perret and Michiels 2021). The purpose for this is felt to be twofold. First, it allows for the redistribution of nuclear proteins such as Sam68 (McBride et al. 1996), polypyrimidine tract binding protein (Back et al. 2002), and the La protein (Meerovitch et al. 1993) to the cytoplasm, where they participate in the translation or replication of viral RNA. Second, it may disrupt nuclear translocation of transcription factors that mediate innate immunity, including interferon response factor 3 (Delhaye et al. 2004) and nuclear factor κB (Watters et al. 2017).

The NPC is a macromolecular structure greater than 100 MDa in size constructed in 8-fold symmetry from subunits known as nucleoporins (Nups). The NPC includes a symmetric core that is comprised of transmembrane Nups that anchor the NPC to the nuclear envelope, inner and outer rings, and a central channel. The central channel consists primarily of phenylalanine-glycine (FG) repeat domains contributed by multiple Nups that confer an intrinsically disordered structure. The central channel creates a permeability barrier against passive diffusion of macromolecules 40 kDa and larger. The central channel also permits active nucleocytoplasmic transport (NCT). NCT is mediated by transport adapters known as importins and exportins which interact with FG-repeats and require a Ran-GTP/Ran-GDP gradient across the nuclear envelope. On the nuclear face of the NPC resides the nuclear basket, which has roles in chromatin organization and RNA quality control, while the cytoplasmic face is adorned with cytoplasmic filaments that are important for transport receptor docking and remodeling of ribonucleoproteins during their export. (reviewed in Lin and Hoelz 2019)

Disruptions of nuclear import by picornaviruses were first reported in cells infected with poliovirus (PV) and coxsackievirus B3 (CV-B3) (Belov et al. 2000). These deficits were later correlated to cleavage of Nup153 and Nup62 during infection with PV (Gustin and Sarnow 2001) or human rhinovirus 14 (RV14) (Gustin and Sarnow 2002), implicating viral proteases in disruption of the NPC. PV 2A protease (2A^pro^) was shown to disrupt the permeability barrier of the NPC, allowing passive diffusion across the nuclear lamina (Belov et al. 2004). RV 3C and 3CD proteases were found to impair nuclear import (Ghildyal et al. 2009). There have been conflicting reports as to whether or not PV 2A^pro^ activity also impairs RNA export (Park et al. 2008, Castello et al. 2009). The list of Nups targeted by enterovirus proteases was expanded to include Nup98 by PV 2A^pro^ (Park et al. 2008) and RanBP2, Nup214, and Nup153 by RV 3C^pro^ and 3CD^pro^ (Ghildyal et al. 2009). A further study in PV-infected cells demonstrated cleavage of Nup98, Nup153, TPR, Nup96, Nup58, Nup35, RanBP2, Nup214, and POM121, without attributing cleavage events to specific proteases (Krull et al. 2010). The cleavage of Nup62 (Park et al. 2010) and Nup98 (Park et al. 2015) by RV2 were found to selectively remove the FG-repeat domain, signifying a potential mechanism for the disruption of transport functions or of the NPC permeability barrier.

Interestingly, though all picornaviruses studied to date alter the NPC, the specific Nup targets and functional outcomes vary between strains. Amongst several RV strains, there were differential kinetics for the degradation of Nup98, Nup153, and Nup62 by 2A^pro^. Most strikingly, some strains showed little or no cleavage of Nup62 while it was efficiently degraded by others (Watters and Palmenberg 2011). Further, the consequences of Nup degradation vary amongst RV strains, showing differential effects on trafficking mediated by various transport receptors including importin-αβ, transportin 1, transportin 3, or Crm1/exportin 1 (Watters et al. 2017). This suggests that the impact of infection by specific picornavirus strains have unique effects on which cargoes are mislocalized, and on the relative balance of import vs export deficits. These differences may have implications for the efficiency of viral replication as well as the potential deleterious effects on host cells.

Few studies have attempted to comprehensively survey the impact of a picornavirus infection on the NPC. A proteomic analysis in cells overexpressing CVB3 2A^pro^ identified several interacting partners of 2A^pro^ and showed that eukaryotic translation initiation factor 4G (eIF4G) and Nup98 were the most rapidly cleaved host proteins. This was associated with bidirectional deficits of nucleocytoplasmic transport and redistribution of nuclear mRNA into cytoplasmic stress granules (Serganov et al. 2022).

Disruption of the nuclear pore complex has been implicated in degenerative disorders of the motor neuron. In *C9orf72*-related amyotrophic lateral sclerosis (ALS), three independent screens identified deficits of nucleocytoplasmic transport contributing to neurodegeneration (Freibaum et al. 2015, Jovicic et al. 2015, Zhang et al. 2015). These results have since been extended to sporadic ALS (Coyne et al. 2021) and other neurodegenerative disorders including Huntington, Alzheimer, and Parkinson diseases (Grima et al. 2017, Eftekharzadeh et al. 2018, Shani et al. 2019). Furthermore, mutations in multiple Nup genes are sufficient to cause neurodegeneration, including several motor neuron diseases of childhood (Juhlen and Fahrenkrog 2018).

Given the well-established role of NPC dysfunction in both picornavirus infection and neurotoxicity, and the similarity of disrupted nucleocytoplasmic transport dysfunction in these settings, we wondered whether NPC dysfunction may be a mechanistic link between EV-D68 infection and neurotoxicity. As a first step in this line of inquiry, we sought to comprehensively characterize the impact of EV-D68 proteases on NPC composition and function, and to determine the contribution of EV-D68 proteases to motor neuron toxicity in models of AFM. Here, we show that 2A^pro^ is primarily responsible for wide-ranging deficits of nucleocytoplasmic transport of protein cargoes via cleavage of a small number of Nups, and is neurotoxic in induced pluripotent stem cell (iPSC)-derived human spinal motor neurons.

## RESULTS

### Enterovirus D68 proteases disrupt nuclear pore complex composition

We screened for disruption of the nuclear pore complex by expressing in HEK293T cells the 2A^pro^ and 3C^pro^ proteases from US/MO/2014-18947, a well-characterized neuropathogenic strain of EV-D68 that has been studied in animal and cell culture models of AFM (Liu et al. 2015, Brown et al. 2018, Hixon et al. 2019). We measured the levels of 30 nucleoporins in whole cell lysates by Western blot. As positive controls, we used the homologous proteases from poliovirus 1 Mahoney strain. To confirm protease activity, we also measured the levels of EIF4G (Ventoso et al. 1998) and CREB (Yalamanchili et al. 1997), well-established non-Nup substrates of 2A^pro^ and 3C^pro^, respectively. After correction for multiple comparisons, we identified 14 Nups whose levels were decreased after expression of EV-D68 2A^pro^ and four Nups whose levels were decreased after expression of 3C^pro^, revealing a widespread disruption of NPC composition following EV-D68 protease expression. In addition, Western blots showed putative cleavage products of three Nups for each protease that were detectable only in the presence of protease (Figure 1A-B). Including these proteins, our total list of candidate Nup substrates of EV-D68 proteases numbered 15 for 2A^pro^ (RanBP2, Nup214, Nup88, Aladin, Gle1, Nup43, Nup96, Nup188, Nup35, POM121, Nup62, Nup54, Nup98, TPR, Nup153) and 6 for 3C^pro^ (RanBP2, Nup214, Nup188, Nup35, NDC1, Nup50).

**Figure 1.**
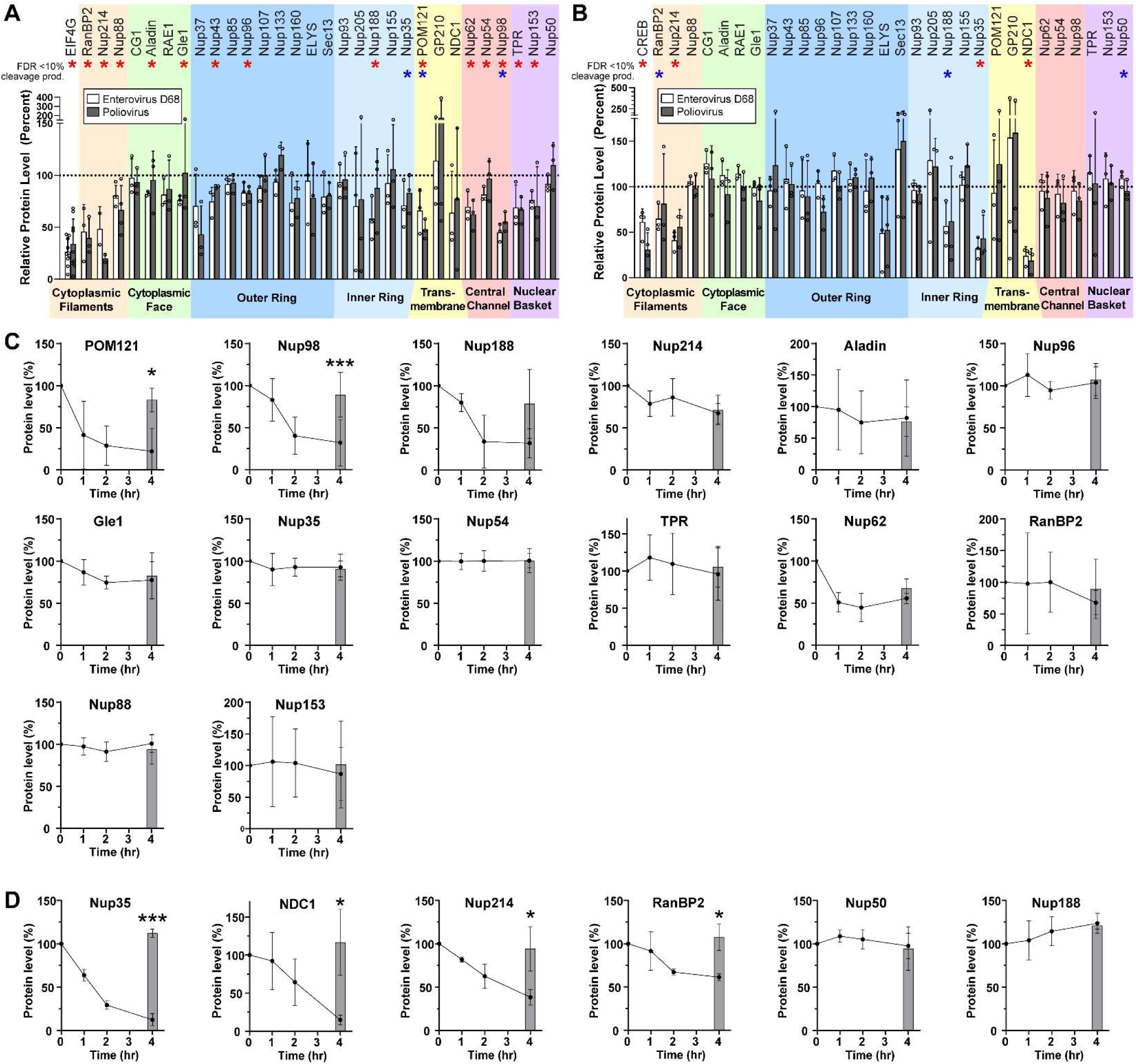
Enterovirus D68 proteases cleave several nucleoporins. (A) HEK cells were transfected with Enterovirus D68 2A^pro^, Poliovirus 1 2A^pro^, or empty vector control, and the indicated nucleoporins were quantified by western blot, normalized to control=100%. EIF4G was included as a non-nucleoporin positive control. Mean ± S.D. of 3 independent replicates. Statistical comparison was by multiple t-tests of the EV-D68 2Apro vs control, with multiple comparisons correction by the false discovery rate method. Red asterisks represent significance with FDR<10%, blue asterisks represent samples for which a cleavage product was visible on the Western blot irrespective of significance. (B) As in A, except transfecting with 3C^pro^. The positive control was CREB. (C) Nuclear lysates from HEK cells were incubated at 37°C with recombinant EV-D68 2A^pro^ at a ratio of 1:50 (protease:lysate by mass) for the indicated time. Grey bars represent lysate incubated for 4 hours at 37°C in the absence of protease as a negative control. Mean ± S.D. of 3 independent replicates. Statistical comparison was by t-test between the with- and without-protease samples at 4 hours. (D) As in C, except with EV-D68 3C^pro^ at a ratio of 1:200. \**p*<0.05, \*\**p*<0.01, \*\*\**p*<0.001.

We considered that alterations in the level of each Nup may be either a direct result of proteolytic cleavage, or a secondary effect such as loss of a binding partner in the complex. We therefore sought to determine which of the candidate Nups are substrates of their respective EV-D68 protease. We performed *in vitro* assays of proteolytic cleavage for each Nup of interest by incubating HEK293T cell nuclear-fractionated lysates with recombinant 2A^pro^ or 3C^pro^ and quantifying Nup levels over time by Western blot. All candidate Nups were detectable by this method with the exception of Nup43, which was excluded from further analysis. Surprisingly, of the 14 remaining candidate substrates of 2A^pro^, only Nup98 and POM121 were significantly proteolyzed in these nuclear preparations (Figure 1C). For 3C^pro^, four out of six candidates demonstrated significant evidence of proteolysis, including Nup35, NDC1, Nup214, and RanBP2 (Figure 1D).

### 2A protease alters nucleocytoplasmic trafficking of protein substrates

We next sought to understand the functional impacts of Nup cleavage and altered NPC composition on nucleocytoplasmic transport. We began by screening for steady state alterations of nucleocytoplasmic localization using a reporter system based on the previously reported Shuttle-tdTomato (Zhang et al. 2015). We expressed NLS-tdTomato, tdTomato-NES, or Shuttle-tdTomato (which we will refer to henceforth as NLS-tdTomato-NES) in HeLa cells, followed by GFP-labeled versions of each of the EV-D68 proteases or an empty vector GFP control. NLS-tdTomato is typically localized to the nucleus and tdTomato-NES to the cytoplasm, serving as readouts of nuclear import and export, respectively. NLS-tdTomato-NES is transported bidirectionally with an approximately uniform nucleocytoplasmic distribution at steady state. Its localization is expected to change during altered nucleocytoplasmic transport only when there are unequal effects on nuclear import vs export. GFP-2A^pro^-expressing cells demonstrated significant alterations, wherein the NLS-tdTomato and tdTomato-NES reporters were significantly less partitioned into the nucleus or cytoplasm (Figure 2A-C; Supplementary Figure 2).

**Figure 2.**
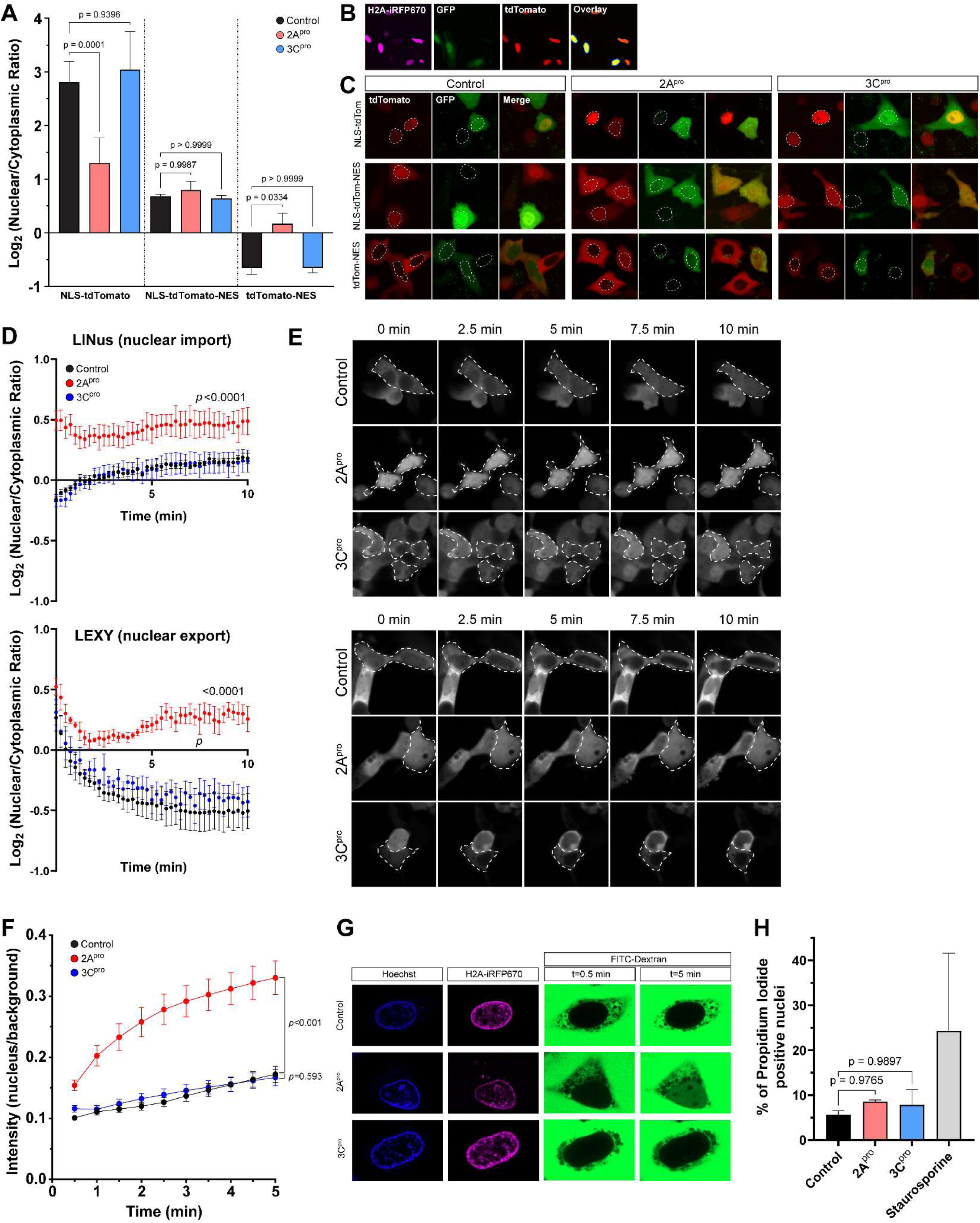
Enterovirus D68 2A protease impairs nucleocytoplasmic transport of protein cargoes and disrupts the nuclear pore complex permeability barrier. (A) tdTomato reporter assay showing nuclear-cytoplasmic ratios at steady state 20 hours after transfection of IRES-GFP, IRES-GFP-2A^pro^, or IRES-GFP-3C^pro^. Mean values of three independent replicates *n*=29,404 GFP+ cells in total, error bars are S.D. Statistical comparison was by two-way ANOVA with Šídák’s multiple comparisons test. (B) Example output of image analysis protocol. In the overlay image, yellow represents the nuclei of transfected cells, defined by the area of expression of H2A-iRFP670 in GFP+ cells only. The surrounding dark blue ring is the perinuclear cytoplasm. The intensity of reporter construct was measured in the red channel and reported as a ratio of nuclear to perinuclear cytoplasmic signal. (C) Example images from A. Dotted outlines mark the nuclei. (D) Kinetics of nuclear import (left) and export (right) measured by photoinducible LINus and LEXY reporter assays, respectively, in HeLa cells 20 hours after transfection with IRES-GFP, IRES-GFP-2A^pro^, or IRES-GFP-3C^pro^. Mean values of 3 biologically independent replicates, *n*=999 GFP+ cells for LINus and 1059 for LEXY in total. Error bars are S.E.M. Statistical comparison was by two-way ANOVA. (E) Example images from D. Dotted outlines mark GFP+ cells. (F) Dextran exclusion assay was performed in HeLa cells 15 hours after transfection with IRES-iRFP670-H2A, IRES-iRFP670-H2A-(2A)-2A^pro^, or IRES- iRFP670-H2A-(2A)-3C^pro^. *n*=45-46 cells per group pooled from 3 biologically independent replicates. Statistical comparison was by two-way ANOVA. (H) Propidium iodide viability assay. Mean values of 3 independent replicates, *n*=8042 cells in total. Error bars are S.D. Statistical comparison was by one-way ANOVA with Tukey’s multiple comparison test.

The above findings have two potential explanations which are not mutually exclusive: (1) 2A^pro^ impairs receptor-mediated nuclear import and export, or (2) 2A^pro^ disrupts the permeability barrier of the NPC, allowing for passive diffusion of macromolecules across the nuclear envelope. To test the first of these possibilities, we utilized a photoinducible reporter system to measure the kinetics of nucleocytoplasmic transport. We used pDB22 (mCherry-LINus) for nuclear import (Niopek et al. 2014) and pDN122 (NLS-mCherry-LEXY) for nuclear export (Niopek et al. 2016). Both mCherry-LINus and NLS-mCherry-LEXY failed to translocate in HeLa cells following expression of 2A^pro^, but not 3C^pro^ (Figure 2D-E; Supplementary Figure 3). These findings demonstrate deficits in both nuclear import and nuclear export of protein cargoes induced by 2A^pro^.

### 2A protease disrupts the permeability barrier of the nuclear pore complex

To assess the integrity of the nuclear permeability barrier we measured the exclusion of a high molecular weight fluorescently labeled dextran from the nucleus (Lenart and Ellenberg 2006) following selective permeabilization of the plasma membrane with digitonin (Adam et al. 1990). As expected, in digitonin-permeabilized HeLa cells expressing an empty vector control or 3C^pro^, FITC-dextran diffused rapidly into the cytoplasm but was excluded from the nucleus. By contrast, in 2A^pro^-expressing cells FITC-dextran accumulated in the nucleus, indicating a disruption of the permeability barrier against diffusion of macromolecules (Figure 2F-G).

In sum, the above data indicate that disruption of the nuclear pore complex specifically by 2A^pro^ leads to deficits in bidirectional nucleocytoplasmic transport of protein cargoes as well as disruption of the NPC permeability barrier against passive diffusion. To exclude the possibility that these phenomena are simply non-specific sequelae of cell death, we measured the uptake of propidium iodide in GFP+ HeLa cells transfected with IRES-GFP, IRES-GFP-2A^pro^, or IRES-GFP-3C^pro^, imaged and analyzed identically to those in the tdTomato and LINus/LEXY reporter assays, and found no significant increase in nuclear propidium iodide uptake between control and protease-expressing cells (Figure 2H).

### Enterovirus D68 proteases do not disrupt RNA export

Given the substantial disruption of nucleocytoplasmic transport of protein cargoes by 2A^pro^, we next asked whether RNA export from the nucleus is similarly altered. We used two complementary measurements of RNA transport in HEK293T cells transfected with EV-D68 proteases: Poly-A RNA-FISH to label the steady-state distribution of native mRNA, and 5-ethinyl uridine (EU) pulse chase labeling of nascent RNAs to measure the kinetics of RNA export. In the case of the latter experiment, we also used combinations of RNA polymerase inhibitors actinomycin D, α-amanitin, and CAS 577784-91-9 to selectively label the products of RNA polymerase I, II, or III. In all cases, there was no significant alteration of nuclear-cytoplasmic localization of RNA (Figure 3A-C; Supplementary Figure 4). We also performed the EU pulse-chase assay in RD cells infected with EV-D68. To control for potential strain-specific effects, we used EV-D68 strains from two different subclades and outbreak years, US/MO/2014-18947 (clade B1, year 2014) and US/MD/2018-23209 (clade B3, year 2018) (Figure 3D-E).

**Figure 3.**
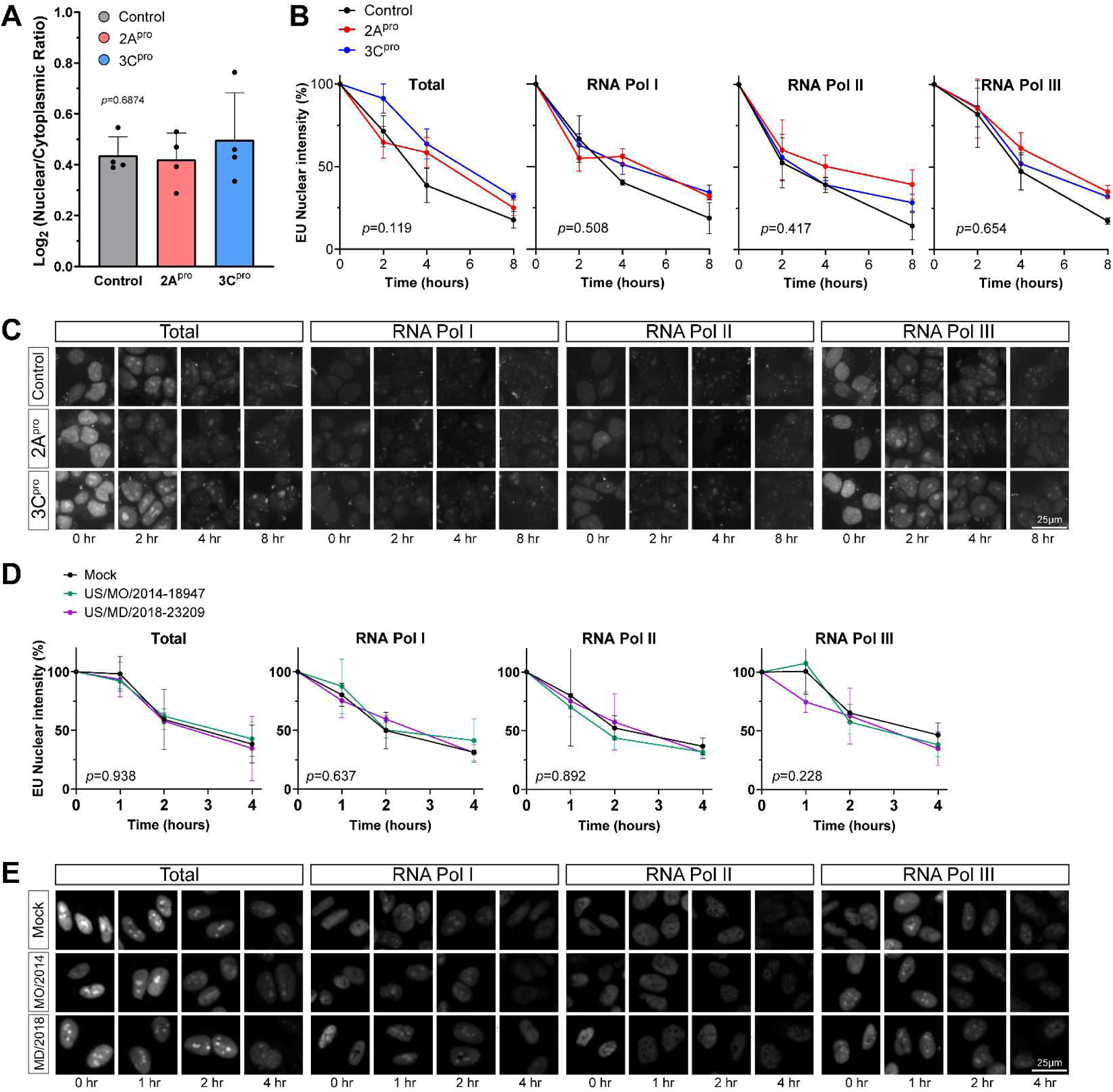
Enterovirus D68 proteases do not significantly alter RNA export. (A) Steady state mRNA levels were measured by Poly-A FISH in HeLa cells 20 hours after transfection with IRES-GFP, IRES-GFP-2A^pro^, or IRES-GFP-3C^pro^. Data represent the mean ± S.D. of four independent replicates, *n*=55,177 GFP+ cells in total. Statistical comparison was by one-way ANOVA with Šídák’s multiple comparisons test. (B) Nuclear export of EU-labeled RNAs in HEK cells by 1hr pulse followed by 0-8hr chase. EU labeling began 20hr after transfection with IRES-GFP, IRES-GFP-2A^pro^, or IRES-GFP-3C^pro^. Pulse labeling of total RNA or primarily the products of the listed RNA polymerases was performed in the presence of pairs of combinations of actinomycin D, α-amanitin, and CAS 577784-91-9 as described in Materials and Methods. Data represent the mean ± S.E.M. of three independent replicates. *n*=253,015 GFP+ cells analyzed in total. Statistical comparison was by two-way ANOVA. (C) Representative images from the experiment described in B. (D) Nuclear export of EU-labeled RNAs in RD cells by 1hr pulse followed by 0-4hr chase. EU labeling began 24 hours after infections with EV-D68 strains at MOI 5 or mock infection as indicated. Data represent the mean ± S.E.M. of three independent replicates. *n*=125,401 cells analyzed in total. Statistical comparison was by two-way ANOVA. (E) Representative images from the experiment described in D.

### Enterovirus D68 infection recapitulates the effect of 2A protease on nucleocytoplasmic transport

We next sought to determine whether key deficits seen during transfection of recombinant proteases were also present during live virus infection. We also focused most of these experiments on a disease-relevant cell type, human iPSC-derived spinal motor neurons (diMNs), using multiple independent iPSC lines to control for the potential variability of these cells. We found that Nup98 and POM121 levels were decreased by EV-D68 infection in a similar manner as 2A^pro^-transfected cell lines, and this was at least partially reversed by the 2A^pro^ inhibitor telaprevir in a dose-dependent manner (Figure 4A). We also evaluated whether nucleocytoplasmic transport disruption occurs during live virus infection by performing the NLS-tdTomato and tdTomato-NES reporter assays in RD cells infected with EV-D68. These cells demonstrated disrupted localization of both markers, and the deficits were rescued in a concentration-dependent fashion by telaprevir. (Figure 4B-C) In diMNs, there was a strong disruption NLS-tdTomato localization in response infection but little if any alteration of tdTomato-NES localization in diMNs. The mislocalization of NLS-tdTomato was partially rescued by 3µM telaprevir. (Figure 4D-E)

**Figure 4.**
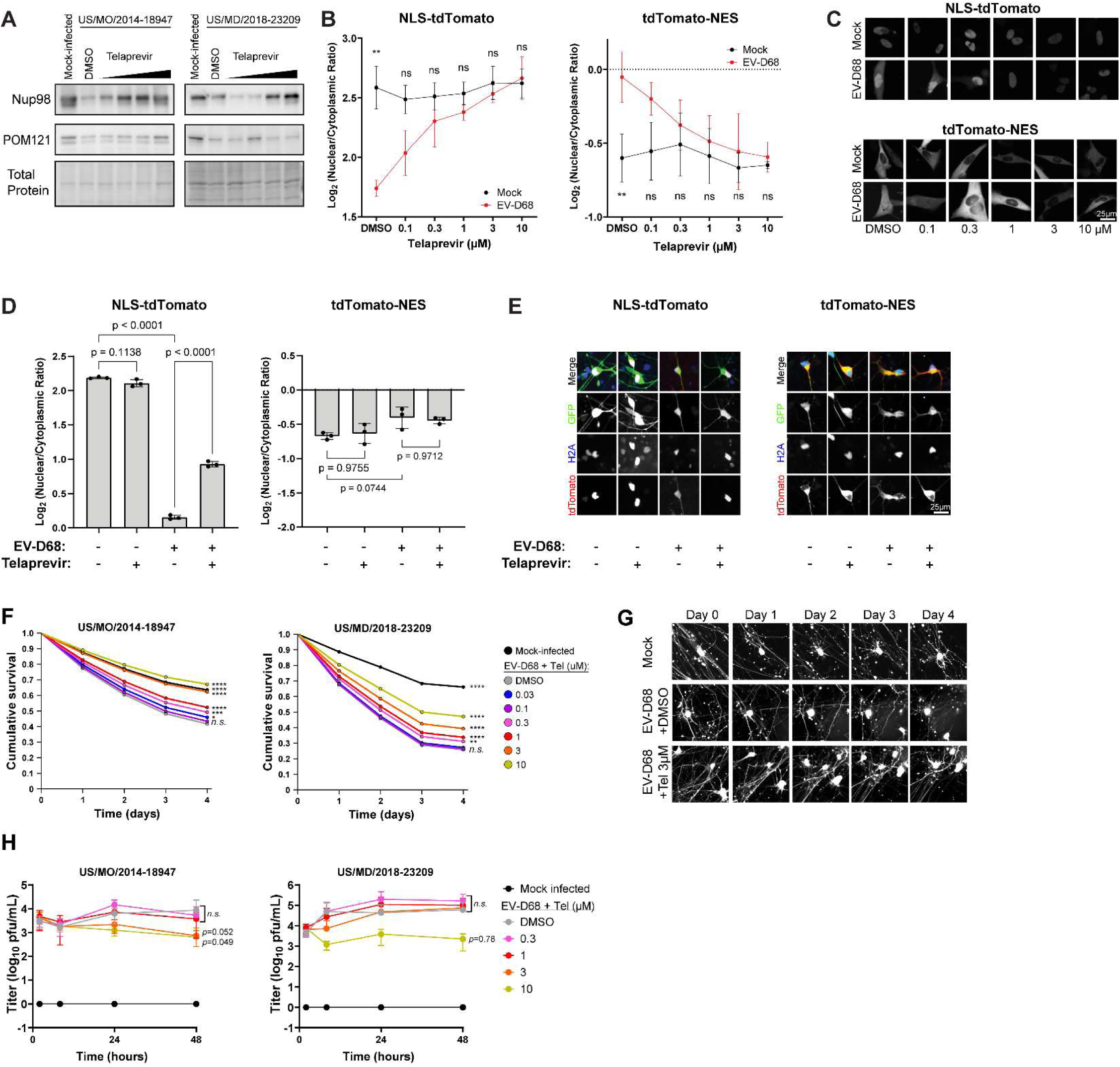
Effects of 2A protease during Enterovirus D68 infection on NPC function and motor neuron toxicity. (A) Western blots of diMN lysates collected 24 hours after infection with the indicated strains of EV-D68 at MOI 5 followed by treatment with telaprevir at 0.3, 1, 3, and 10 µM. Total protein loading control was Biorad stainfree imaging. One representative example out of three biologically independent replicates. (B) tdTomato reporter assays in RD cells showing nuclear-cytoplasmic ratios 24 hours after mock infection or infection with EV-D68 US/MO/2014-18947. Telaprevir was added following infection at the indicated concentrations. Data are mean of three independent replicates, *n*=11,847 cells in total for NLS-tdTomato and *n*=8941 for tdTomato-NES. (C) Representative images of tdTomato signal from B. (D) tdTomato reporter assays in diMN showing nuclear-cytoplasmic ratios 24 hours after mock infection or infection with EV-D68 US/MO/2014-18947, treated with DMSO or 3µM telaprevir. Means of three replicates from independent iPSC lines. *n*=3388 cells in total for NLS-tdTomato and. *n*=3371 for tdTomato-NES. (E) Representative images from panel D. (F) Survival of diMNs following infection with EV-D68 at MOI 5 with the indicated EV-D68 strains in the presence of varying concentrations of telaprevir or DMSO control. diMNs were generated from four independent iPSC lines. *n=*8512 and 7046 cells counted for US/MO/2014-18947 and US/MD/2018-23209 respectively. Statistical comparisons were by Cox proportional hazard regression. (G) Representative images of diMNs from the experiment presented in panel F. Note fragmentation of neurites and loss of cell body shape signifying cell death following EV-D68 infection, and relative preservation following treatment with 3µM telaprevir. (H) Growth curve of EV-D68 in the presence of varying concentrations of telaprevir or DMSO control. Infections were performed with the indicated strains at MOI 0.5, using diMNs generated from four independent iPSC lines for each virus. Error bars are SEM. Statistical comparisons were by two-way ANOVA with Dunnet’s multiple comparisons test between DMSO and telaprevir-treated groups. \**p<*0.05, \*\**p<*0.01, \*\*\**p<*0.001, \*\*\*\**p<*0.0001.

### Inhibition of 2A protease rescues the toxicity of Enterovirus D68 to spinal motor neurons out of proportion to its potential antiviral effect

In the above experiments, we demonstrated that EV-D68 2A^pro^ disrupts the composition of the nuclear pore complex and has deleterious consequences on the maintenance of the NPC permeability barrier and nuclear transport of protein cargoes in cell lines and human spinal motor neurons. We next sought to determine the extent to which 2A^pro^ activity independently contributes to motor neuron injury during EV-D68 infection. GFP-labeled diMNs were infected with EV-D68 and imaged daily by high content imaging with automated cell counting. These diMNs demonstrated gradual yet robust cell death in culture. The EV-D68 2A^pro^ inhibitor telaprevir led to a dose-dependent reduction in cell death, with statistically significant neuroprotection occurring at concentrations of 0.3µM and above (Figure 4F-G). We considered the possibility that the rescue of nucleocytoplasmic transport deficits and motor neuron toxicity may simply be due to an antiviral effect of telaprevir. However, the growth curve of EV-D68 in diMNs showed modest inhibition by telaprevir that achieved statistical significance in only one of two strains at a concentration of 10µM, and in neither strain at lower concentrations (Figure 4H). We therefore infer that telaprevir protects motor neurons from EV-D68-induced cell death primarily because it inhibits the effects of 2A^pro^ activity on host targets.

## DISCUSSION

Here we have presented a detailed evaluation of the effects of EV-D68 proteases on the composition and function of the NPC. Our findings on Nup cleavage are compatible with previous findings in other picornaviruses, and extend the literature by presenting one of the most comprehensive evaluations of Nups to date. Interestingly, we found widespread disruption of the NPC by EV-D68 proteases. The levels of up to 16 Nups were decreased by one or both proteases. By contrast, only 6 Nups were found to be direct proteolytic substrates of 2A^pro^ or 3C^pro^. This finding suggests that the remaining Nup alterations may be a secondary effect. For example, some of the altered Nups may be anchored to the NPC by other Nups which are themselves targets for proteolysis.

Our study also provides a mechanistic explanation for disrupted localization of nucleocytoplasmic cargoes. By using several complementary reporter systems to detect abnormalities of nucleocytoplasmic transport dynamically and at steady state we showed that the deficits are multifactorial, resulting from both disruption of the NPC’s permeability barrier against passive diffusion of macromolecules and from loss of receptor-mediated nuclear import and export.

It is striking that functional deficits in the NPC were entirely attributable to 2A^pro^, even though we only observed two definitive substrates of 2A^pro^: Nup98 and POM121. Nup98 has been well-studied as a target of picornavirus 2A proteases, though studies have drawn differing conclusions regarding the extent and nature of the impact of Nup98 cleavage on NPC function (Park et al. 2008, Castello et al. 2009, Serganov et al. 2022). The finding that 2A^pro^ also cleaves POM121 is especially interesting because of its role in motor neuron degeneration in ALS. POM121 appears to function as a “keystone” for the NPC. In models of ALS, removal of POM121 from the NPC is sufficient to cause loss of multiple Nups including Nup98, Nup107, Nup50, TPR, Nup133, GP210, and NDC1 (Coyne et al. 2020), a list that overlaps with what we observed following 2A^pro^ expression. In these ALS models, overexpression of POM121 is sufficient to reverse NPC deficits, and the mechanism of POM121 removal appears to be inappropriate activation of the ESCRT III pathway (Coyne et al. 2021, Coyne and Rothstein 2021, Baskerville et al. 2024). In the context of EV-D68 infection, it remains to be determined whether Nup98 or POM121 is the primary driver of NPC pathology, if the mechanism of POM121 loss involves the ESCRT pathway or is only due to its cleavage by 2A^pro^, and whether deficits of NPC function can be reversed via repair of Nups or prevention of their cleavage. These questions will be a focus of future studies.

We have also demonstrated that 2A^pro^ activity contributes to nucleocytoplasmic transport dysfunction and separately to cell death in motor neurons infected with EV-D68. Our approach was motivated by the fact that similar NPC deficits have been shown to be toxic in neurodegenerative disorders of the motor neuron. We therefore hypothesize that the 2A^pro^-dependent toxicity that we witnessed in diMNs is at least partially due to these deficits of NPC function. Concordant with our observation of 2A^pro^-dependent toxicity, a recent study demonstrated amelioration of the AFM phenotype in EV-D68-infected mice treated with telaprevir. The protection of spinal motor neurons in these mice occurred despite minimal effects on viral replication in the spinal cord, providing further support for a model in which telaprevir’s mechanism of action is primarily neuroprotective rather than antiviral (Frost et al. 2023). A significant limitation of our study, however, is that we cannot exclude potentially toxic effects of 2A^pro^ on aspects of host neuronal biology aside from the NPC. For example, 2A^pro^ impairs 5’-cap-dependent RNA translation via eIF4G cleavage (Ventoso et al. 1998) and inhibits stress granule formation (Visser et al. 2019) which could also contribute to toxicity in EV-D68-infected neurons. Further work will be required to disentangle the relative contributions of these pathways to motor neuron injury.

## CONCLUSION

We have built upon prior observations in several picornaviruses to show that EV-D68 proteases cause broad disruption of the composition and function of the nuclear pore complex, and that these phenomena are attributable to the cleavage of a relatively small number of Nups by 2A^pro^. Further, 2A^pro^ contributes to motor neuron death in a model of EV-D68 infection. These findings motivate future work to evaluate the NPC or 2A^pro^ as potential therapeutic targets to treat or prevent paralysis in AFM.

## MATERIALS AND METHODS

### Key Reagents

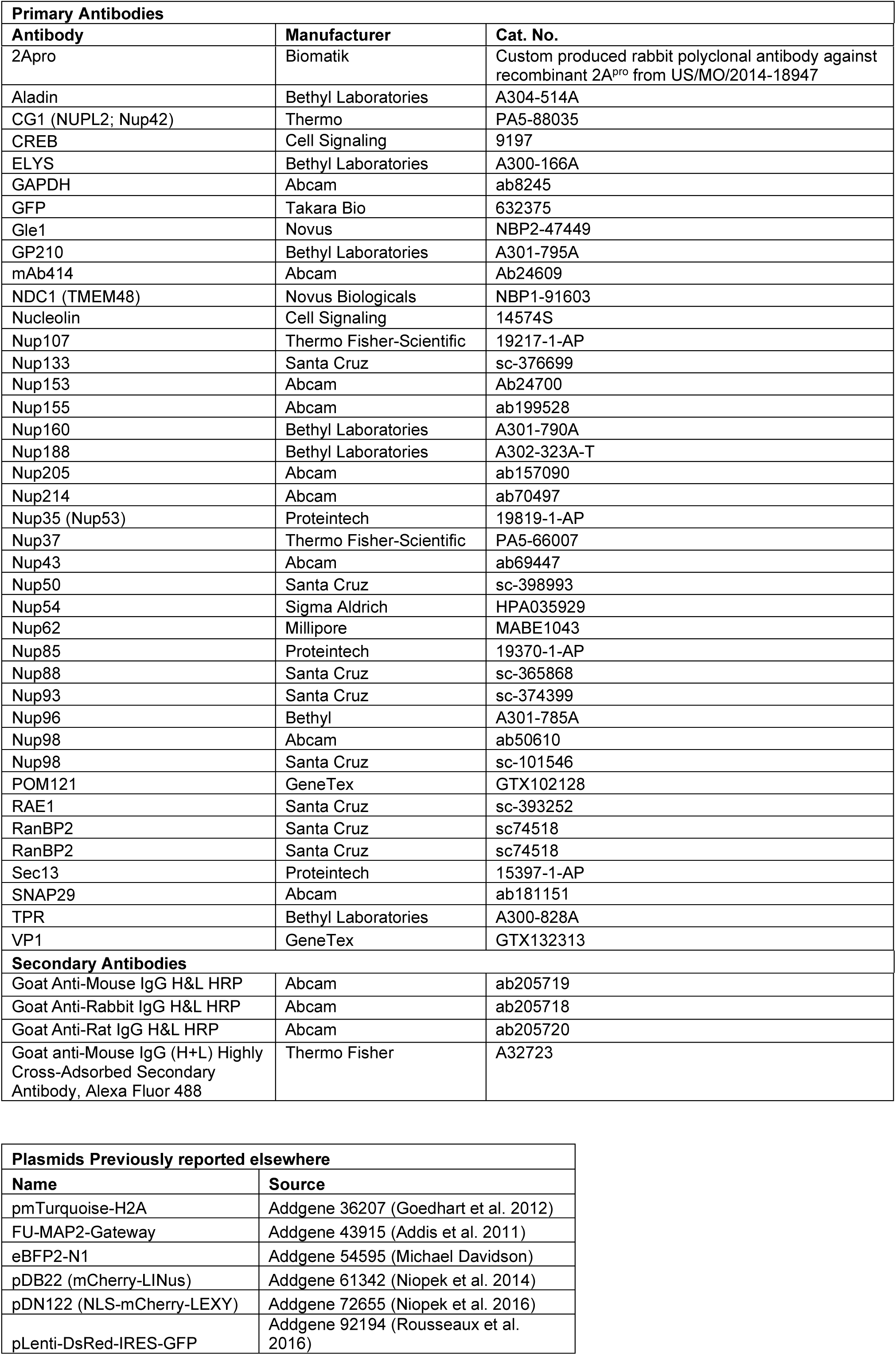
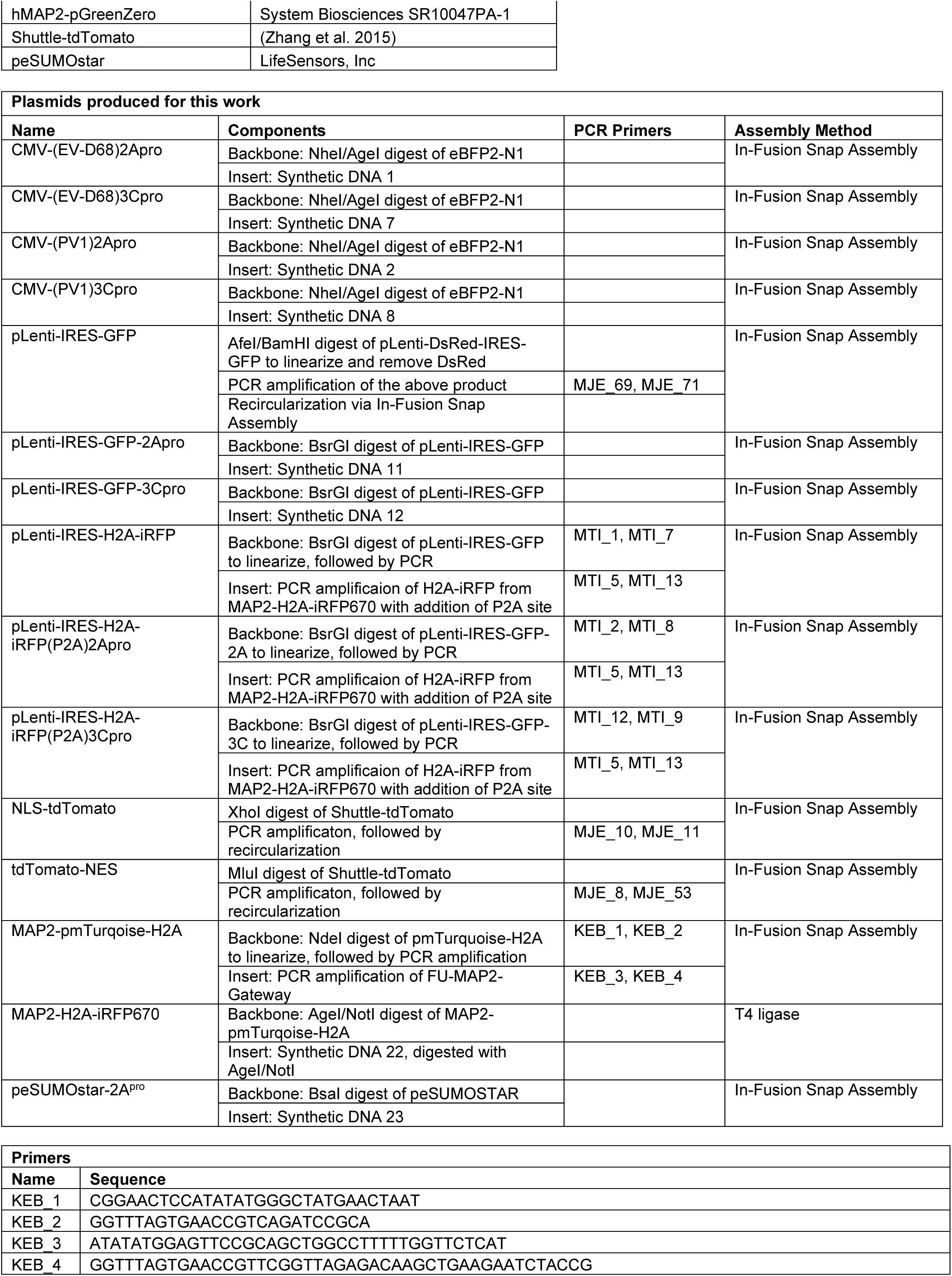
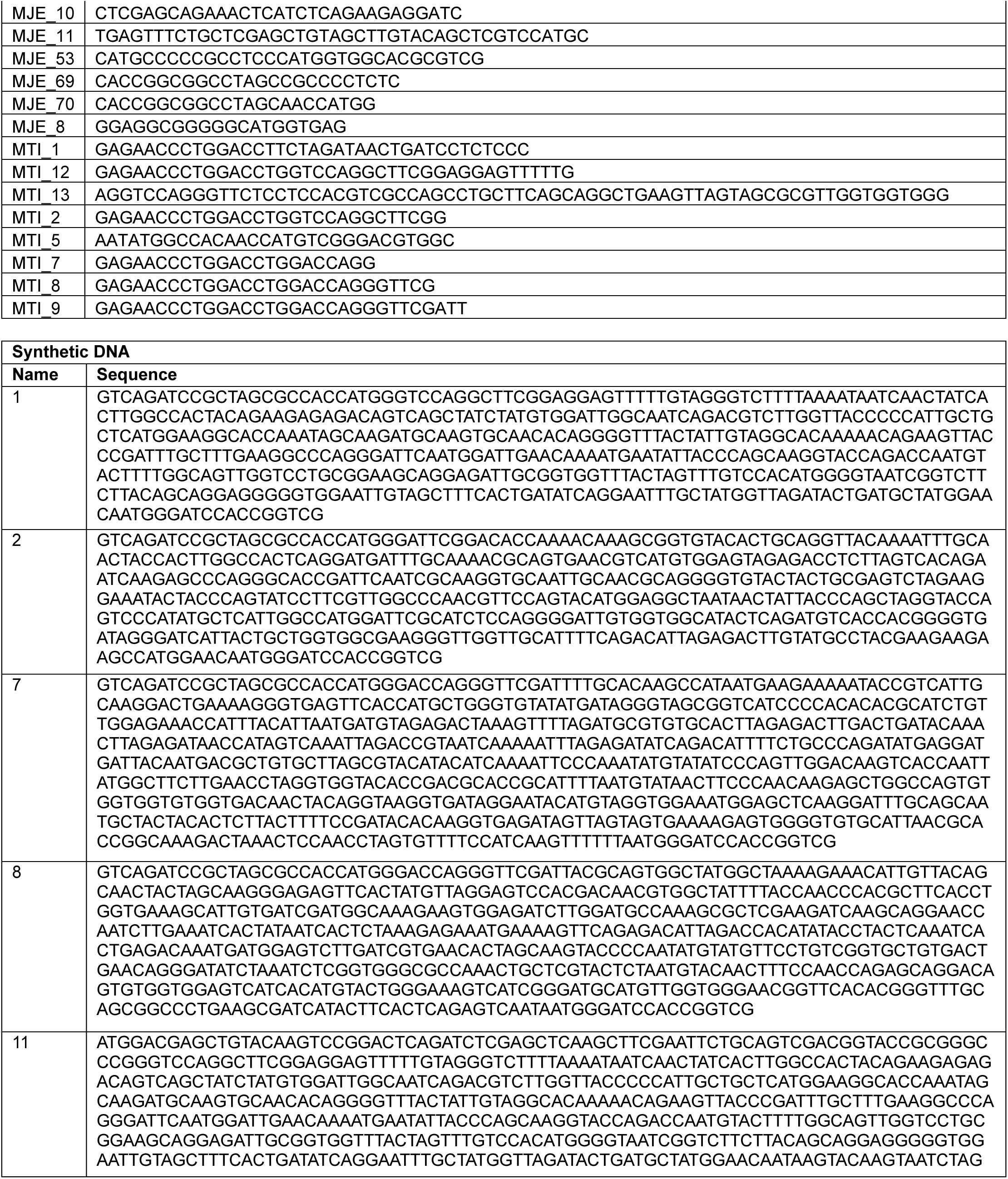
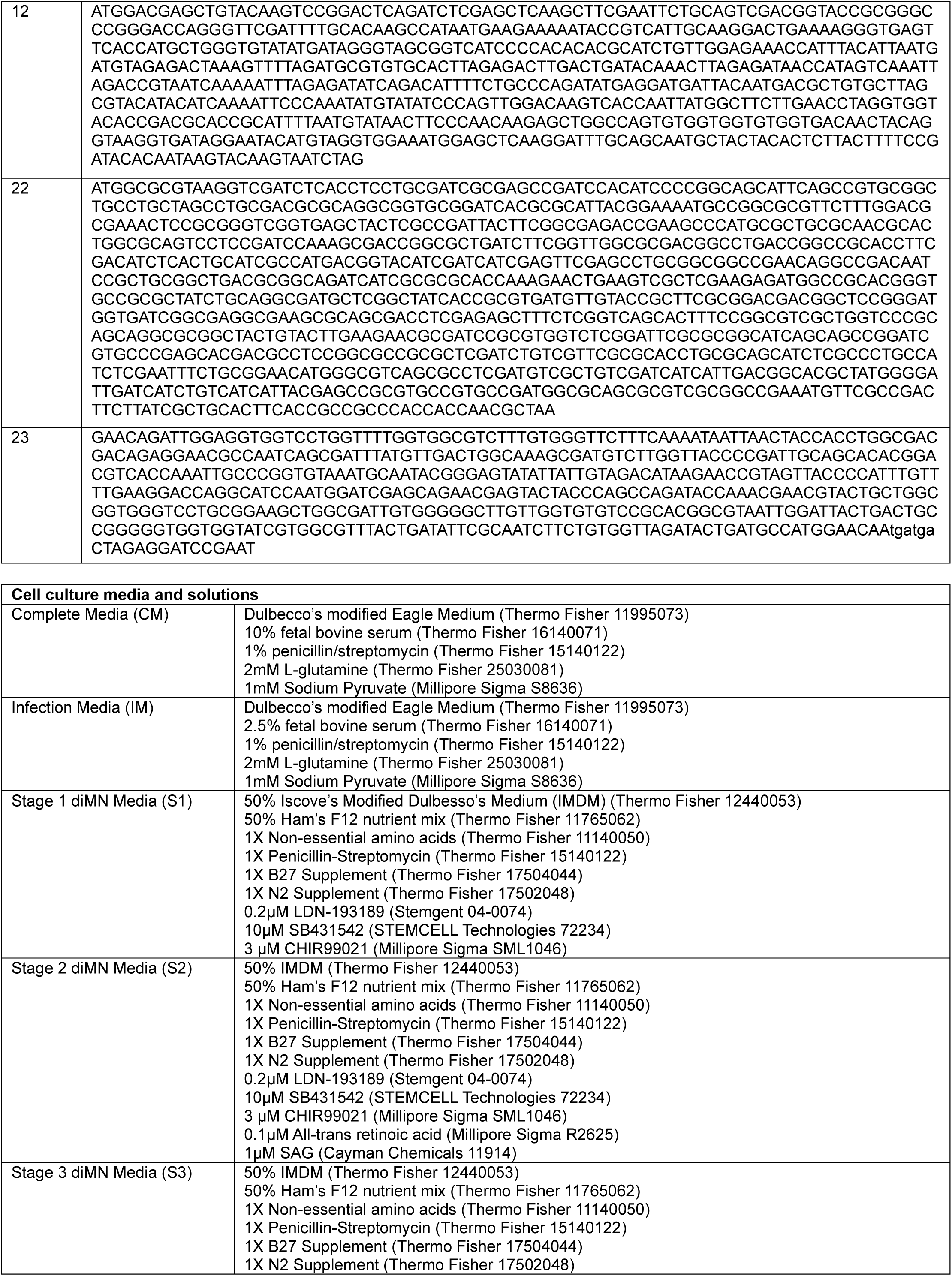
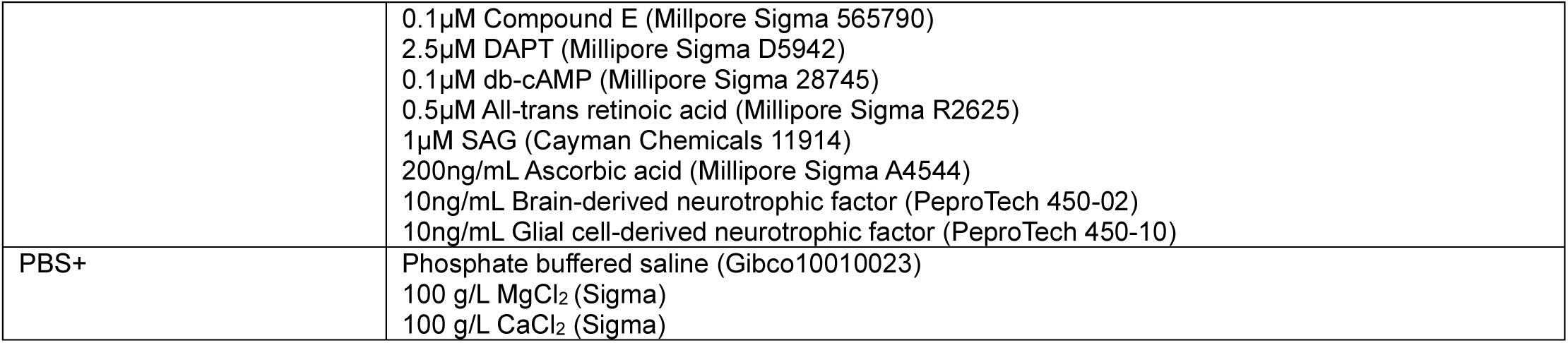

### Plasmid Construction

Plasmids were constructed from the components listed in the Key Reagents table using In-Fusion Snap Assembly (Takara Bio) or T4 DNA Ligase (New England Biolabs) following the manufacturers’ protocol. The products of restriction digests and PCR reactions were purified using 1% agarose gel electrophoresis. The band of interest was visualized using SYBR Safe dye (Thermo Fisher) under blue light to avoid UV exposure, cut out of the gel, and recovered using the GeneJet Gel Extraction Kit (Thermo Fisher). Transfections of non-neuronal cell lines were performed using FuGene HD per the manufacturer’s protocol (Promega). Transfection of motor neurons was by Lonza Nucleofection as described below. Of note, expression of EV-D68 proteases using a CMV promoter and a Kozak sequence as the ribosome binding site, while sufficient to introduce measurable protease activity (Figure S1), did not allow a sufficiently bright signal for imaging assays using fluorescently tagged proteases. This is a commonly encountered problem, owing to the inhibition of transcription and 5’-cap-dependent translation by enterovirus proteases, which we resolved by incorporating an internal ribosome entry site (IRES) in front of the fusion protein (Castello et al. 2006).

### Viruses and titering

Viral strains were US/MO/2014-18947 (BEI Resources, NR-49129) and US/MD/2018-23209 (gift of Dr. Andrew Pekosz, Johns Hopkins Bloomberg School of Public Health). Viral stocks were prepared by inoculating RD cells cultured to approximately 90% confluency in T75 flasks at 33°C and 5% CO_2_ at an MOI of 0.01 in Infection Media (IM). Once cytopathic effect was evident (3-5 days) lysates were collected by three freeze-thaw cycles. Titering was performed by methylcellulose plaque assay. RD cells were plated in 6-well tissue culture plates and grown to confluency. A ten-fold dilution series of each viral sample was prepared in IM. RD cells were washed with PBS+ and the viral dilutions were added in a volume of 250µl/well and then placed in the 33°C incubator for one hour with gentle rocking every 15 minutes. The viral inoculum was aspirated, 2mL of 1% methylcellulose in MEM was added to each well as an overlay medium, and the plates were returned to the 33°C incubator for 3 days. To fix and harvest the plates, 2 mL of 10% neutral buffered formalin (VWR, 10015-194) was added dropwise and incubated overnight at room temperature. The formalin/cellulose mixture was swirled and decanted off, and then the cells were stained with naphthol blue black (Millipore Sigma) overnight and washed with water. Plaques were counted manually in a well containing 30-100 plaques, and pfu/ml was calculated by (number of plaques) / (0.25 mL) x (dilution factor).

### Cell lines and induced pluripotent stem cells

RD cells, HEK293T cells, and HeLa cells were from ATCC (CCL-136, CRL-3216, and CRM-CCL-2, respectively). iPSC lines were from the Cedars Sinai Biomanufacturing Center repository, and included lines CS0002, CS0003, CS88, CS0YX7, and CS8VTR. iPSCs were maintained in Essential 8 Flex media (Thermo Fisher) on 6-well tissue culture plates coated with Cultrex UltiMatrix Reduced Growth Factor Basement Membrane Extract (R&D Systems BME001), manually cleaned to remove differentiated cells, and passaged as colonies using ReLeSR passaging reagent (STEMCELL Technologies). Motor neuron differentiation was performed following the direct-induced motor neurons (diMN) protocol reported previously (NeuroLINCS Consortium et al. 2021). In brief, iPSCs were grown to 30-40% confluency and then fed daily with S1 media for 6 days for neuroepithelial induction. On day 6, neuroepithelial cells were collected following incubation with Accutase (Millipore Sigma), using gentle pipetting to release cells from the centers of the colonies. Neuroepithelial cells were triturated to a single cell suspension and then collected by centrifugation at 200*g*. These cells were cryopreserved using Cryostor (STEMCELL Technologies) for storage in liquid nitrogen until later use. Day 6 neuroepithelial cells were thawed and replated into UltiMatrix-coated tissue culture dishes in S2 media with ROCK inhibitor Y-27632 (Cayman Chemical) at 20µM, and then fed daily on days 7-12 with S2 without ROCK inhibitor. This stage generates motor neuron precursor cells. On day 12, motor neuron precursors were collected as a single cell suspension with 0.05% Trypsin and replated onto the desired format for each experiment. Where indicated, nucleofection was performed at this step, prior to replating. From day 12 onward, cells were fed with S3 media to promote terminal maturation and maintenance of motor neurons. The diMN protocol produces cultures containing approximately 80% Tuj1^+^ neurons, including 40% Islet-1^+^ mature motor neurons and 40% Nkx6.1^+^ immature motor neurons (NeuroLINCS Consortium et al. 2021).

### Western blotting

Cells were harvested by scraping in ice-cold PBS, pelleted by centrifugation at 500*g* for 5 minutes at 4°C, and PBS was aspirated off. Lysis was achieved by resuspension in ice-cold RIPA buffer (Millipore Sigma) supplemented with cOmplete protease inhibitor cocktail (Millipore Sigma) followed by sonication. Protein concentrations were measured by DC protein assay (Bio-Rad). Samples were prepared by diluting a constant protein quantity by mass with deionized water and 4X Laemmli buffer (Bio-Rad) to a final concentration of 1X. Protein electrophoresis was performed in Bio-Rad Criterion Stain-Free gels and then transferred onto low-fluorescence PVDF membranes using the Transblot Turbo system (Bio-Rad). Total protein stain for confirmation of proper transfer and normalization of quantitative data was by Ponceau S stain (Bio-Rad; Figures 1A-B) or by stain-free imaging (Figures 1C-D, S1). Blots were blocked in 5% non-fat milk in TBS with 0.1% Tween-20 (TBS-T) for 1 hour. Primary antibodies were applied in TBS-T overnight at 4°C. Blots were then washed three times in TBS-T and incubated for one hour in secondary antibody at a concentration of 1:20,000. Blots were washed three more times with TBS-T, twice with TBS, and then developed using SuperSignal West Atto ECL substrate (Thermo Fisher). Images were acquired using a GE Healthcare ImageQuant LAS 4000 (Figures 1A-B) or Bio-Rad ChemiDoc MP Imaging System (Figures 1C-D, S1). Quantification of western blot images was performed using ImageJ software and normalized to the total protein stain.

### Recombinant Enterovirus D68 proteases

Recombinant proteases were produced based on the sequence of US/MO/2014-18947. 3C^pro^ was produced by Biomatik. 2A^pro^ produced by standard methods did not demonstrate evidence of protease activity. We therefore generated a SUMO-2A^pro^ fusion protein, based on a strategy previously reported to circumvent this technical limitation thought to be due to improper folding (Musharrafieh et al. 2019). The sequence of 2A^pro^ was codon-optimized for E. coli using the IDT codon optimization tool (https://www.idtdna.com/pages/tools/codon-optimization-tool) and generated as synthetic DNA (Integrated DNA Technologies) for In-Fusion Snap Assembly into the peSUMOstar plasmid as detailed above to encode His_6_-SUMO-2A^pro^. BL21(DE3) competent cells (New England BioLabs) were transformed with peSUMOstar-2A^pro^, grown in a shaker-incubator at 37°C in the presence of kanamycin 50µg/mL to an optical density of 0.6-0.75, cooled to 18°C, and then protein expression was induced with 0.5mM IPTG for 24 hours. Cells were harvested by centrifugation at 3500*g* for 20 minutes. The cell pellet with resuspended and lysed in 50mM sodium phosphate, 300 mM sodium chloride, 10nM imidazole, 1mg/mL lysozyme (Roche), and 3 units/mL Benzonase nuclease (Millipore Sigma) at pH 8.0 and incubated on ice for 30 minutes. The lysate was loaded onto Ni-NTA spin columns (Qiagen) and washed with a buffer containing 50mM sodium phosphate, 300 mM sodium chloride, and 20nM imidazole at pH 8.0, then eluted with a buffer containing 50mM sodium phosphate, 300 mM sodium chloride, and 500nM imidazole at pH 8.0.

### Recombinant protease digestion assay

To generate a substrate enriched for nucleoporins, a nuclear lysate was collected from HEK293T cells using the NE-PER extraction kit (Thermo Fisher). Nuclear lysates and recombinant proteases were dialyzed overnight at 4°C into reaction buffer (20 mM HEPES pH 7.4, 150 mM potassium acetate, and 1 mM DTT). Lysates were diluted to a concentration of 1µg/µL in reaction buffer with proteases added at a ratio (protease:lysate by mass) of 1:50 for SUMO-2A^pro^ and 1:200 for 3C^pro^. Proteases were omitted in the case of negative control samples. The samples were incubated at 37°C with rotation at 300rpm for the indicated times. The reaction was stopped via the addition of 4X Laemmli buffer to a final concentration of 1X, followed by incubation for 5 minutes at 100°C. Samples were analyzed by Western blotting.

### Nucleocytoplasmic transport assays

HeLa cells or RD cells were plated onto UltiMatrix-coated 96-well glass-bottom plates (Cellvis), cultured in Complete Media, and transfected on day 1 with a reporter construct (NLS-tdTomato, NLS-tdTomato-NES, tdTomato-NES, mCherry-LINus, NLS-mCherry-LEXY) and the nuclear marker H2A-iRFP670. On day 2, they were transfected with the protease-containing vector or empty vector control, or infected with EV-D68 mock-infected. Imaging was completed at the indicated number of hours after the second transfection or infection on a Molecular Devices ImageXpress Confocal high content imaging system. For photoinducible reporters (mCherry-LINus and NLS-mCherry-LEXY), images were acquired every 15 seconds, each beginning with a 1-second pulse of the 488nm laser at maximum intensity to maintain the reporter in its activated state. Image analysis was performed in automated fashion using MetaXpress software (Molecular Devices). A custom analysis protocol was developed to select only GFP^+^ cells with size and morphology typical of live cells, to define the area of the nucleus using the H2A-iRFP670 signal, exclude nuclei with a condensed or mitotic morphology, and to define the perinuclear cytoplasm as a concentric ring outside of the nucleus. The average intensity of the reporter signal in the nucleus and perinuclear cytoplasm were measured and reported as Log_2_ of the nucleus/cytoplasm ratio. Cells were excluded from the analysis when the intensity of the reporter signal was too low to be distinguished from background or high enough to create shine-through artefact in the green channel, up to a maximum of 50% of the pixel depth of the image to ensure signal in the linear range. Due to inherent variability in signal intensity, these minimum and maximum intensity values were determined empirically for each independent biological replicate.

For assays in diMN, cells were nucleofected with hMAP2-pGreenZero, MAP2-H2A-iRFP670, and the reporter construct on day 12 of differentiation and plated to UltiMatrix-coated 96-well glass-bottom plates at a density of 2.5×10^5^ cells/well. On day 18, they were infected with EV-D68 at an MOI of 2.5. Imaging was performed at 24 hours post-infection. Analysis was as above, except that the perinuclear cytoplasm was defined as the intersection between the cytoplasmic GFP signal and a concentric ring outside the nucleus.

### Dextran exclusion assay

Digitonin permeabilization was carried out as previously described (Hayes et al. 2020) with some adjustments. Briefly, HeLa cells were seeded onto a UltiMatrix-coated 96-well glass-bottom plate (Cellvis) and maintained in Complete Media. Cells were transfected 15 hours before imaging at 60-70% confluency with bicistronic reporter constructs encoding a nuclear marker and the protease of interest or empty vector (pLenti-IRES-H2A-iRFP, pLenti-IRES-H2A-iRFP(P2A)2Apro, or pLenti-IRES-H2A-iRFP(P2A)3Cpro). To permeabilize, cells were kept on ice and washed with ice-cold PBS for 5 minutes. Digitonin was added at varying concentrations for 10 minutes in permeabilization buffer (PB; 20 mM HEPES, 110 mM KOAc, 5 mM Mg(OAc)2, 0.5 mM EGTA, 250 mM sucrose) to identify the concentration that permeabilized the plasma membrane while leaving the nuclear envelope intact. The optimized digitonin concentration varied depending on the cell density, but the same concentration was used across each biological replicate, with 30-40 µg/mL being used for analysis. Permeabilized cells were washed twice for 5 minutes with modified transport buffer (TRB; 20 mM HEPES, 110 mM KOAc, 2 mM Mg(OAc)2, 5 mM NaOAc, 0.5 mM EGTA, 250 mM sucrose). Cells were kept in TRB with Hoechst (5 µg/mL) on ice until imaging.

Imaging was performed on a Zeiss LSM880 Airyscan confocal microscope and processed using ImageJ. FITC-conjugated 70kDa dextran (Millipore Sigma) was diluted in TRB and added to each well immediately before imaging at a final concentration of 0.6 mg/mL. Images were acquired every 30 seconds for 5 minutes. For each field, cells were analyzed only if the nuclei expressed H2A-iRFP670, had intact and visible nuclear membranes, were regularly shaped, and had clear and sharp boundaries between the nucleus and the cytoplasm. Images were excluded if a majority of the cells in the field were permeabilized regardless of transfection status, indicating general over-permeabilization. Cells were also excluded if they were impermeabilized (indicated by the absence of a visible nuclear membrane), if nuclei were not completely within the plane of imaging, or if there was drift of the focal plane during time course imaging. Data were reported as the average FITC intensity in the nucleus divided by the average FITC intensity of the extracellular background.

### Propidium iodide viability assay

HeLa cells were transfected as in the nucleocytoplasmic transport assays, except omitting the reporter construct. Staurosporine 2 µM was added to the positive control group two hours before propidium iodide as a positive control. Propidium iodide 100µg/mL was added to all groups 20 hours post-transfection and 15 minutes prior to imaging. Image acquisition and analysis was the same as for the nucleocytoplasmic transport assays, selecting only GFP+ cells with normal size and morphology. Instead of calculating the nuclear-cytoplasmic ratio of signal in the red channel, the intensity of signal in the nucleus was recorded. To determine the minimum intensity cutoff for propidium iodide positivity, histograms were plotted for signal intensity in the staurosporine-treated group, and a cutoff for positive cells was determined by identifying the inflection point in the biphasic distribution.

### Poly-A FISH

HeLa cells were plated on UltiMatrix-coated 96-well glass-bottom plates (Cellvis) and transfected with pLenti-IRES-GFP, pLenti-IRES-GFP-2A^pro^, or pLenti-IRES-GFP-3C^pro^. Twenty hours later, they were washed with PBS, fixed for 30 minutes in 4% paraformaldehyde, washed two more times with PBS, then incubated 10 min each with ice-cold 100% methanol followed by 100% ethanol. Cells were incubated with 1ng/µL of Cy3-oligo-dT (Genelink) in hybridization buffer (30µM sodium citrate, 300µM sodium chloride, 10% formamide, 10% dextran sulfate, 50µg yeast tRNA (Millipore Sigma) in DEPC-treated water for 1 hour at 37°C. To enhance the GFP signal, the cells were then immunofluorescently stained for GFP. Cells were washed with PBS and then blocked in blocking solution (5% goat serum, 1% bovine serum albumin in PBS) for one hour at room temperature and then incubated in primary antibody against GFP diluted 1:200 in 1/4x blocking solution overnight at 4°C. The cells were then washed twice with PBS and then incubated with AlexaFluor488-conjugated secondary antibody for 1 hour in 1/4x blocking solution. Finally, the cells were washed with PBS, counterstained with Hoechst 33342 nuclear stain (Cell Signaling) at 5µg/mL for 15 minutes, and then mounted with PBS glycerol for imaging. Imaging was performed and analyzed on a Molecular Devices ImageXpress Confocal high content imaging system as described above for NCT assays.

### 5-Ethinyl Uridine pulse-chase

HEK293T cells were plated on UltiMatrix-coated 96-well glass-bottom plates (Cellvis) and transfected with pLenti-IRES-GFP, pLenti-IRES-GFP-2A^pro^, or pLenti-IRES-GFP-3C^pro^. Twenty hours after transfection, the cell culture medium was replaced with fresh media containing 1mM 5-ethinyl uridine (EU; Vector Laboratories) for a 1-hour pulse to label nascent RNA. To preferentially label the products of each RNA polymerase, inhibitors of the other two RNA polymerases were included during the EU pulse. These were 0.05ng/mL actinomycin D, 0.05 ng/ml α-amanitin, and 50 µM CAS 577784-91-9 to inhibit RNA polymerases I, II, and III, respectively. The EU-containing media was aspirated off, washed once with PBS, and then chased with fresh Complete Media for 0, 2, 4, or 8 hours. For RD cells, the cells were infected with EV-D68 at MOI5 and the pulse-chase labeling was begun 24 hours after infection, with the chase being completed at 0, 1, 2, or 4 hours after the end of the pulse. Cells were collected by fixation with 3% paraformaldehyde for 15 minutes, washed twice with PBS, and then permeabilized with 0.5% Triton-X100 for 15 minutes. EU was fluorescently labeled with AZDye594-azide via click chemistry using the Click&Go cell reaction kit following the manufacturer’s instructions (Vector Laboratories). The cells were immunofluorescently co-stained for GFP and counterstained with Hoechst 33342 as described above for the poly-A-FISH assay. Imaging and analysis was likewise performed as above, except nuclear signal intensity of the EU stain was reported instead of nuclear/cytoplasmic ratio, because the cytoplasmic signal was not consistently distinguishable from background. Intensities were normalized such that the intensity at t=0 was considered 100%.

### Motor neuron toxicity

Day 12 diMN precursors were transfected with hMAP2-pGreenZero using a Lonza Nucleofector 4D X unit, with the P3 primary cell kit on program DC-104. Cells were replated into UltiMatrix-coated 96-well glass bottom plates at a density of 100,000 cells/well in S3 media with 20µM Y-27632. After 24 hours, the cells were re-fed without Y-27632 and maintained in S3 media until day 32 of differentiation. These diMNs were infected with EV-D68 at MOI 5 in Infection Media for one hour at 37°C with rocking every 15 minutes, then fed with S3 media. diMNs were imaged on a Molecular Devices ImageXpress Confocal high content imaging system immediately following infection (considered t=0) and then once daily for four additional days. GFP^+^ neurons were counted via automated image analysis in MetaXpress software.

### Growth curves on motor neurons

Day 12 diMN precursors were plated onto UltiMatrix-coated 96-well plastic tissue culture plates at a density of 75,000 cells/well and maintained in S3 media until day 32 of differentiation. diMNs were infected with EV-D68 at MOI 0.5 as described above. Virus was collected at the indicated times by subjecting the plates to three freeze-thaw cycles. Viral titers were measured by methylcellulose plaque assay.

### Statistics

Statistical analyses and preparation of graphs were performed in GraphPad Prism 10 software. Statistical methods for each experiment are described in the corresponding figure legend.

## ACKNOWLEDGEMENTS

We thank Dr. Andrew Pekosz for helpful discussions and the gift of US/MD/2018-23209. We thank Lin Jin, Lin Xue, and Kelly Bowen for technical expertise. The following reagent was obtained through BEI Resources, NIAID, NIH: Enterovirus Species D Type 68, US/MO/14-18947 (produced in serum-free A549 cells), NR-52013. This work was supported by the National Institutes of Health (5K 12NS098482, K08 NS124989, R01 NS143998 and P50 HD103538).

## SUPPLEMENTARY DATA

**Figure S1.**
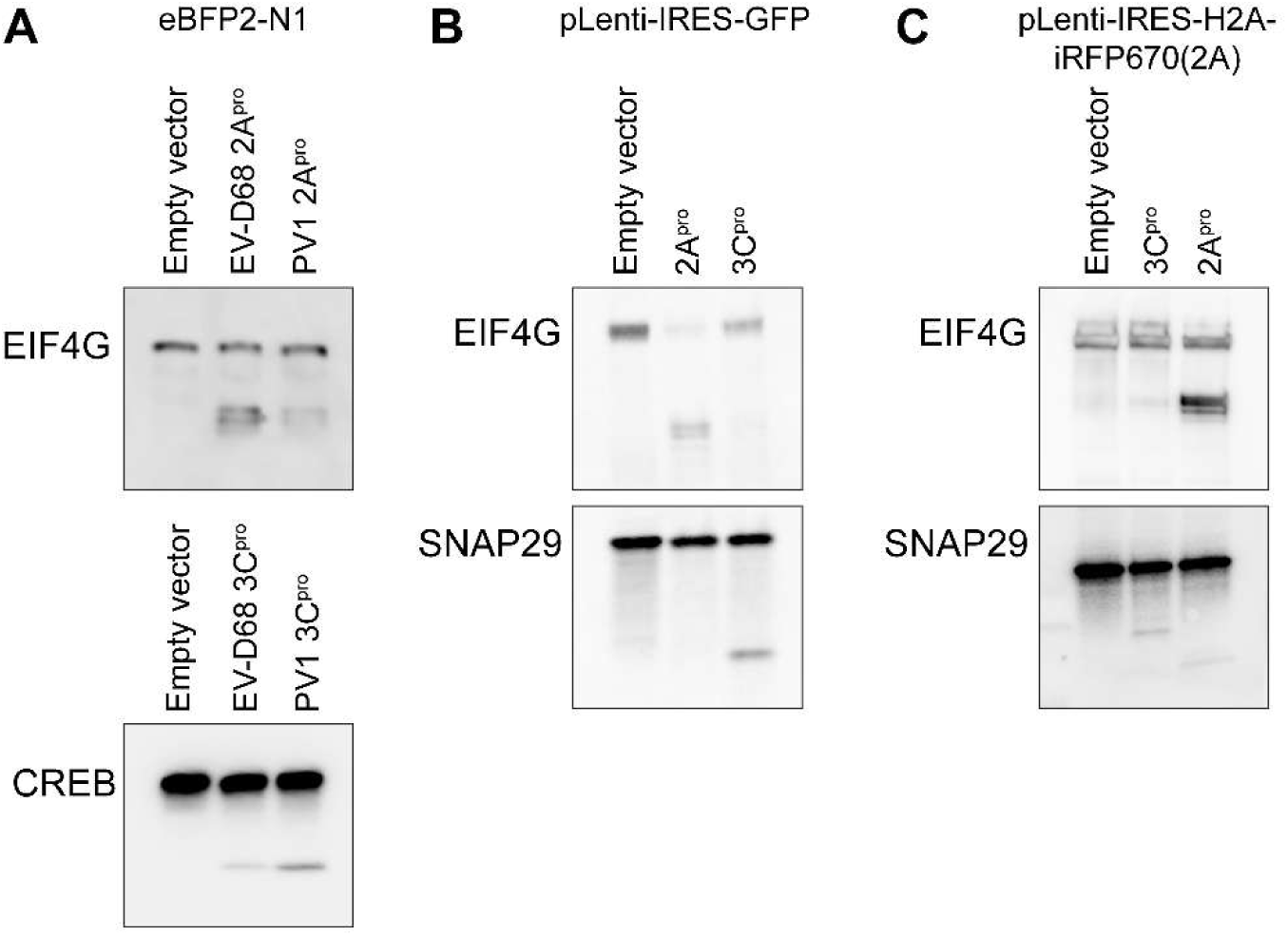
Western blots demonstrating evidence of 2A^pro^ and 3C^pro^ protease activity on well-established substrates following transfection with each of the expression vectors used in this study.

**Figure S2.**
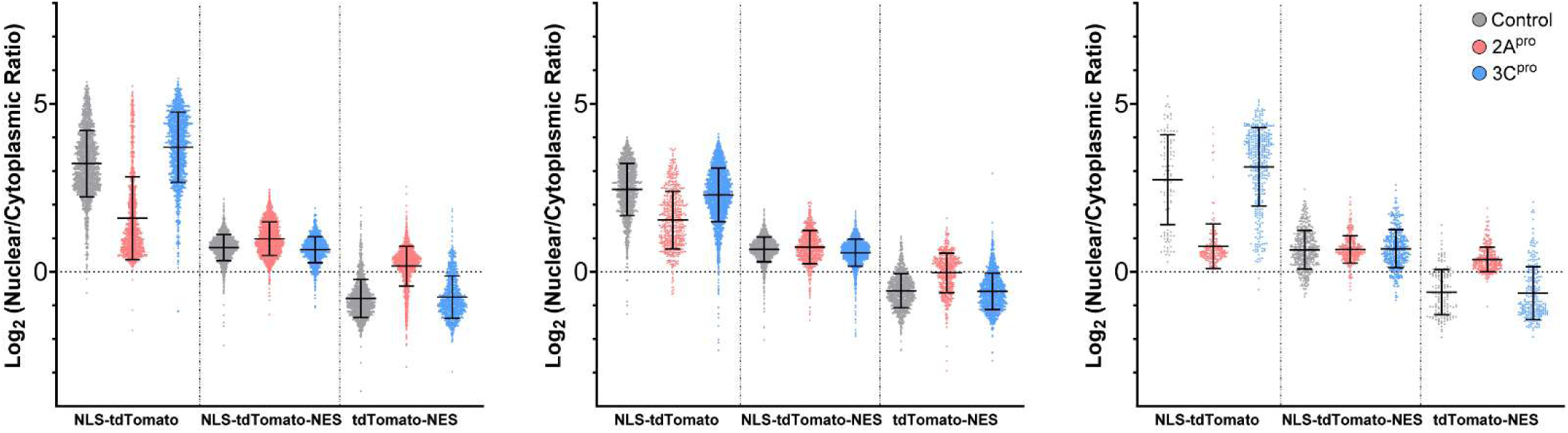
Source data for Figure 2A, showing individual cell level data for three independent replicates. Bars are mean ± S.D.

**Figure S3.**
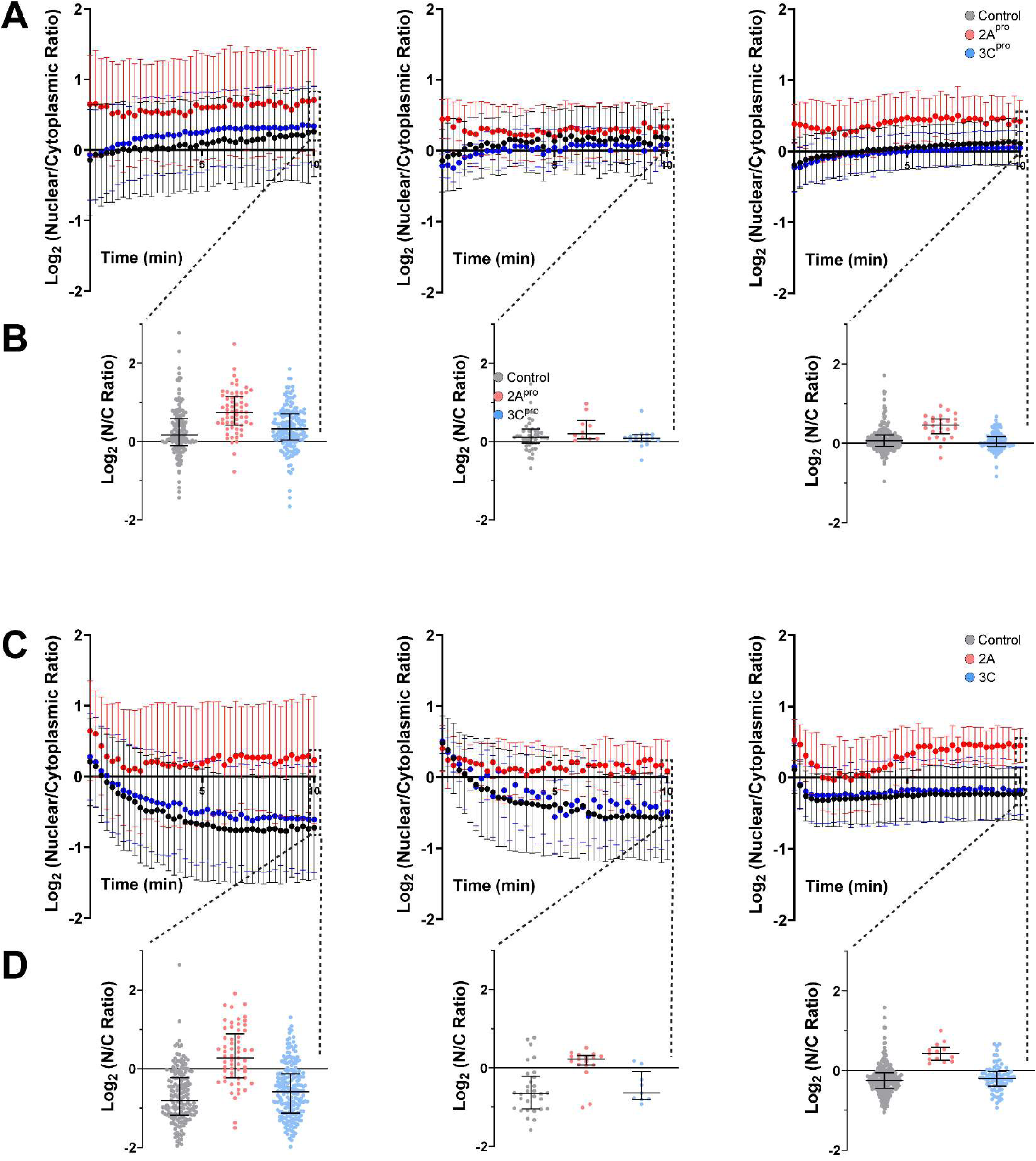
Source data for Figure 2D, showing mean ± S.D. values from three independent replicates for LINus (A) and LEXY (C) nucleocytoplasmic transport reporters. Individual cell level data is also presented at t=10 min for LINus (B) and LEXY (D). Bars are mean ± S.D.

**Figure S4.**
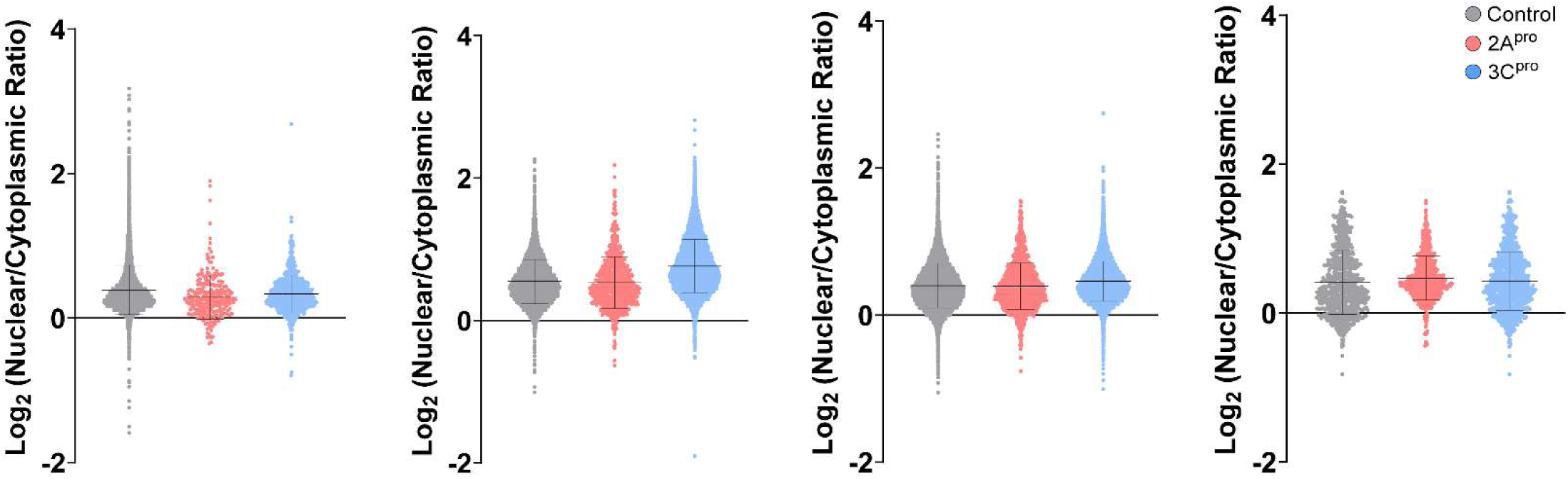
Source data for Figure 3A, demonstrating individual cell-level data across four independent replicates. Bars are mean ± S.D.

**Figure S5.**
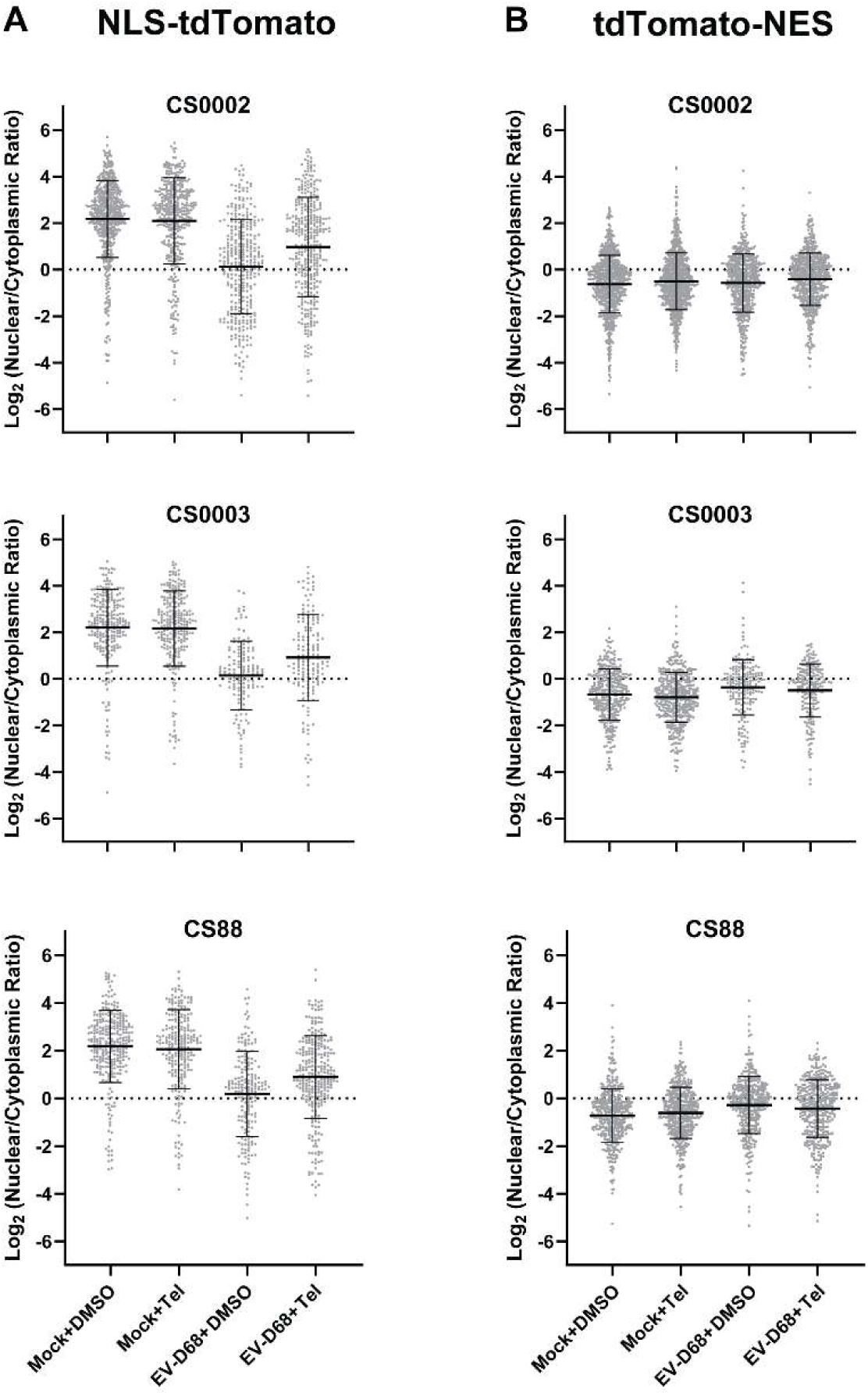
Source data for figures 4C (A) and 4D (B), showing individual cell-level data for three independent replicates in diMNs derived from iPSC lines CS0002, CS0003, and CS88. Bars are mean ± S.D.

